# Neuronal CCL2 expression drives inflammatory monocyte infiltration into the brain during acute virus infection

**DOI:** 10.1101/208587

**Authors:** Charles L Howe, Reghann G. LaFrance-Corey, Emma N Goddery, Kanish Mirchia

## Abstract

**Background:** Viral encephalitis is a dangerous compromise between the need to robustly clear pathogen from the brain and the need to protect neurons from bystander injury. Theiler's murine encephalomyelitis virus (TMEV) infection of C57Bl/6 mice is a model of viral encephalitis in which the compromise results in hippocampal damage and permanent neurological sequelae. We previously identified brain infiltrating inflammatory monocytes as the primary driver of this hippocampal pathology, but the mechanisms involved in recruiting these cells to the brain were unclear.

**Methods:** Chemokine expression levels in the hippocampus were assessed by microarray, ELISA, RT-PCR, and immunofluorescence. Monocyte infiltration during acute TMEV infection was measured by flow cytometry. CCL2 levels were manipulated by immunodepletion and by specific removal from neurons in mice generated by crossing a line expressing the Cre recombinase behind the synapsin promoter to animals with floxed CCL2.

**Results:** Inoculation of the brain with TMEV induced hippocampal production of the proinflammatory chemokine CCL2 that peaked at 6 hours postinfection, whereas inoculation with UV-inactivated TMEV did not elicit this response. Immunofluorescence revealed that hippocampal neurons expressed high levels of CCL2 at this timepoint. Genetic deletion of CCR2 and systemic immunodepletion of CCL2 abrogated or blunted the infiltration of inflammatory monocytes into the brain during acute infection. Specific genetic deletion of CCL2 from neurons reduced serum and hippocampal CCL2 levels and inhibited inflammatory monocyte infiltration into the brain.

**Conclusions:** We conclude that intracranial inoculation with infectious TMEV rapidly induces the expression of CCL2 in neurons, and this cellular source is necessary for CCR2-dependent infiltration of inflammatory monocytes into the brain during the most acute stage of encephalitis. These findings highlight a unique role for neuronal production of chemokines in the initiation of leukocytic infiltration into the infected central nervous system.

## Background

Neural injury associated with virus infection represents a multifaceted convergence of host-pathogen interactions that range from direct lytic killing of infected neurons to bystander pathology mediated by brain-infiltrating immune cells responding to chemotactic and inflammatory cues. Viral encephalitis is a trade-off between the need to clear pathogen from the brain and the need to preserve irreplaceable neurons and neural circuits: too little inflammation and the host dies of uncontrolled infection, too much inflammation and the host suffers permanent brain damage [1]. And while much attention is rightly given to viral encephalitides associated with human mortality, there is likely a significant component of neural injury associated with low-level, sub-clinical viral infections of the central nervous system that are ultimately cleared by the host. Theiler's murine encephalomyelitis virus (TMEV) is a model of such an infection [2]. When C57Bl/6 mice are inoculated via intracranial delivery of the Daniel's strain of TMEV there is an acute viral encephalitis that culminates in generation of an antiviral T cell-mediated response, development of virus neutralizing antibodies, clearance of the virus, and resolution of brain inflammation over the course of about 45 days [3]. However, despite essentially complete resolution of the infection, permanent neurological sequelae such as impaired spatial learning [2, 4], anxiety [5], and epilepsy [6] occur in most post-infectious animals. Uniquely, these neurologic problems largely stem from bystander loss of CA1 pyramidal neurons and subsequent disruption of hippocampal and hippocampal-cortical circuits [4, 7, 8].

Our previous studies identified brain infiltration of inflammatory monocytes, a population defined as CD45^hi^CD11b^+^Gr1^+^1A8^−^ cells, as the primary driver of hippocampal pathology during acute TMEV infection [4, 8]. We showed that animals which mount a large inflammatory monocyte response exhibit extensive loss of CA1 neurons in the dorsal hippocampus and lose the ability to learn spatial navigation and novel object recognition tasks. In contrast, mice that mount a weak inflammatory monocyte response exhibit preservation of CA1 neurons and maintain cognitive performance, despite robust virus infection [8]. In parallel, others have shown that monocyte-derived inflammatory factors such as interleukin-6 [9, 10] and tumor necrosis factor-α [11] drive ictogenesis in the TMEV model. These observations suggest that modulation of inflammatory monocyte responses during acute virus infection in the brain may confer neuroprotection. However, the specific mechanisms responsible for the recruitment of inflammatory monocytes to the brain in the TMEV model have not been previously characterized and open questions remain regarding the mechanisms of leukocyte infiltration into the brain in general.

The multistep process of leukocyte entry into the central nervous system is predominantly controlled by chemokines [12]. In particular, inflammatory monocyte trafficking is thought to depend upon C-C motif chemokine receptor type 2 (CCR2) signaling in response to C-C motif chemokine ligand 2 (CCL2) [13], though the specific details of this dependency vary with viral pathogen. For example, CCR2-deficient mice exhibited reduced monocyte recruitment to the brain but increased mortality during West Nile virus encephalitis [14] and this effect was differentially regulated by CCL2 and CCL7, another CCR2 ligand [15]. In a model of Japanese encephalitis virus infection, mice deficient in CCR2 had reduced monocyte infiltration and reduced mortality, whereas CCL2-deficient animals exhibited increased monocyte infiltration and increased mortality [16]. Both CCL2- and CCR2-deficient mice with mouse hepatitis virus encephalitis had reduced monocyte infiltration, but only CCR2-deficient mice exhibited increased mortality [17, 18]. Likewise, mice with CCR2-deficient hematopoietic cells mounted a reduced monocyte response to herpes simplex virus 1 infection and exhibited increased mortality [19]. These studies clearly support a role for the CCR2:CCL2 axis in trafficking of inflammatory monocytes to the brain during viral encephalitis. However, the specific cellular source of the CCL2 is not identified in any of this work. In the current study we demonstrate that inflammatory monocyte infiltration into the brain during acute TMEV infection requires CCR2 and that neurons are a key source of CCL2 driving this trafficking during the earliest stages of infection.

## Methods

### Mice

C57BL/6J (#000664), Syn-Cre (#003966), and CCL2-RFP^fl/fl^ (#016849) mice were acquired from The Jackson Laboratories (Bar Harbor, ME). Mice were acclimatized for at least one week following shipment and prior to use or were bred in-house. Female mice between 4-6 weeks of age were used for all experiments. LysM:eGFP mice were maintained in-house, as described [4]. Syn-Cre and CCL2-RFP parents were bred in a our facility to establish double homozygous F2 hybrids (Syn-Cre^+/+^CCL2-RFP^fl/fl^). All mice were group housed under controlled temperature and humidity with a 12 hour light/dark cycle with ad libitum access to food and water. All animal experiments were performed according to the National Institutes of Health guidelines and were approved by the Mayo Clinic Institutional Animal Care and Use Committee (Animal Welfare Assurance number A3291-01).

### Virus and infection

At 4-6 weeks of age, mice were infected by intracranial injection of 2×10^5^ PFU of the Daniel’s strain of TMEV in 10 μL DMEM, prepared as previously described [20]. Sham-infected mice received intracranial injection of 10 μL virus-free DMEM (Cellgro; Herndon, VA). In some experiments, virus was inactivated by exposure to short-wavelength UVc light (254 nm) for 15 minutes at room temperature immediately prior to inoculation.

### RNA and microarray

RNA was isolated from the whole brain of mice following perfusion with ice cold PBS using the RNeasy Lipid Tissue Midi Kit (Qiagen, Valencia, CA). RNA integrity, purity, and concentration was assessed using an RNA Analysis kit (Agilent Technologies, Santa Clara, CA). Samples passing quality control were analyzed on Illumina mouse WG-6 v 2.0 expression BeadChips in the Mayo Clinic Medical Genome Facility Gene Expression Core. Expression data were analyzed using Excel and MATLAB, where fold change was calculated and converted to log_2_. Heatmaps and hierarchical clusters were derived using Gitools v2.2.2.

### Cytokine measurements

Following TMEV infection, serum and whole brain or hippocampal homogenates were collected, clarified, and stored at −80°C until analysis. Mouse CCL2 was detected using the Quantikine ELISA kit (R&D Systems, Minneapolis, MN) following the manufacturer’s instructions. For each hippocampus and whole brain sample, 50 μL of neat homogenate was measured in duplicate. Serum samples were diluted 2-fold and measured in duplicate. Values were calculated from a standard curve included in every assay. Serum values are reported as per mL; tissue homogenate values were back-calculated to the total amount present in the entire animal.

### Immunohistochemistry and microscopy

Terminally anesthetized mice (isoflurane overdose) were perfused with 50 mL of 4% paraformaldehyde (PFA) via intracardiac puncture and tissues were postfixed in 4% PFA for 24 hours. Brain tissue was macrosectioned using a brain matrix, making cuts through the optic chiasm and infundibulum. The resulting tissue block containing the dorsal hippocampus was embedded in 4% agarose gel and sectioned at 70 μm thickness by vibratome. Free floating sections were blocked in PBS containing 1% BSA, 10% normal donkey serum, 1% FBS, and 0.1% Triton-X 100 for 1 hour, incubated overnight at 4°C with primary antibody (anti-MCP-1/CCL2: Cell Sciences, CPM001, 1:400 in block), incubated with secondary antibody for 1 hour (donkey anti-rabbit Cy3: Jackson ImmunoResearch, 711-166-152, 1:1000 in block), washed, and mounted in DAPI-containing mountant on charged slides. CCL2 immunoreactivity was imaged with the Zeiss AxioObserver.Z1 and ApoTome.2 structured illumination system (Carl Zeiss Microscopy GmbH, Jena, Germany) using a 20× objective (LD Plan-Neofluar 20×/0.4 Korr Ph 2 M27, 0.55 NA), 538-562 nm bandpass excitation, 570-640 nm bandpass emission, 7 μm optical thickness, and 300 msec exposure time. For detection of the mCherry fluorophore in CCL2:RFP animals, free floating 70 μm sections were mounted on charged slides in DAPI-containing mountant. Fluorescence images were captured with a laser scanning confocal microscope (LSM780, Carl Zeiss Microscopy GmbH, Jena, Germany) using a 40x objective (C-Apochromat 40×/1.20 W Korr FCS M27, 1.2 NA) with water as the refractive medium. Validation of mCherry-specific emission and exclusion of autofluorescence was obtained with spectral imaging using a lambda scan at 488 nm, 561 nm, and 594 nm excitation and 8 nm-stepped emission spectra. 40 μm z-stacks were acquired with a step thickness of 2 μm, pixel dwell time of 0.39 μs, and pinhole equivalent to 1 airy unit across all samples. Uncompressed TIFF-images were exported from Zen software (Zen Black 2012 64-bit, Carl Zeiss Microscopy GmbH, Jena, Germany) and post-processed in ImageJ (ImageJ v1.50b, Wayne Rasband, National Institutes of Health, USA) and Photoshop (Adobe Photoshop CC, 2014 Release, 64-bit). Levels were normalized, when appropriate, equally across images and groups; gamma values were not changed.

### Isolation of brain-infiltrating leukocytes (BILs)

BILs were isolated as previously described [20]. Briefly, following cardiac perfusion with 50 mL PBS, leukocytes were isolated from Dounce homogenized whole brain tissue using a 30% Percoll gradient centrifuged at 7800 g_ave_ for 30 min at RT in a Beckman F0630 rotor. The floating myelin layer was removed and the leukocytes were collected, strained at 40 μm via gravity, diluted in 50 mL PBS, and centrifuged at 600 g for 5 min at RT in a Beckman SX4250 rotor. The leukocyte pellet was resuspended in 1 mL PBS and underlaid with 1 mL of 1.100 g/mL Percoll to enrich for monocytes and neutrophils. With subsequent centrifugation at 800 g for 20 min at RT in a Beckman SX4250 rotor without brake, the mononuclear leukocytes were collected at the gradient interface, washed in PBS, and resuspended in flow cytometry buffer containing 1% bovine serum albumin and 0.02% sodium azide in PBS.

### Flow cytometric phenotyping

BILs were incubated with 0.5 μg Fc block (anti-FcyRIII/II mAb) prepared from the supernatant of 2.4G2 hybridoma cells for 30 min on ice, followed by staining with CD45 (clone 30-F11, BD Biosciences), CD11b (clone M1/70, BD Biosciences), Ly6G/C (clone RB6-8C5, BD Biosciences), or Ly6G (clone 1A8, BD Biosciences). All antibodies were added to blocked wells at 1:200, incubated for 30 min, and washed three times prior to flow cytometric analysis. Brain infiltrating cells were gated on CD45 expression. All CD45^mid/hi^ cells were further assessed for expression of CD11b, Ly-6C/G (Gr1), and Ly-6G (1A8). We defined inflammatory monocytes as the CD45^hi^CD11b^+^Gr1^++^1A8^−^ population, neutrophils as CD45^hi^CD11b^++^Gr1^+^1A8^+^ cells, and microglia as CD45^mid^CD11b^mid^Gr1^−^ cells. For phenotyping experiments involving reporter LysM:eGFP mice, we defined inflammatory monocytes as GFP^mid^ cells, neutrophils as GFP^hi^ cells, and microglia as GFP^neg^ cells within a specific forward-and side-scatter gate. Flow cytometric analysis was performed on an Accuri C6 flow cytometer with sampler arm (BD Biosciences, Mountain View, CA). Files were analyzed offline using FlowJo 10.08 (Windows version; FlowJo LLC, Ashland, OR).

### Statistics and data analysis

α=0.05 and β=0.2 were established a priori. Post hoc power analysis was performed for all experiments and significance was only considered when power ≥0.8. Statistical analyses were performed in JMP Pro 12 (SAS Institute Inc., Cary, NC). Normality was determined by the Shapiro–Wilk test and normally distributed data were checked for equal variance. Parametric tests were only applied to data that were both normally distributed and of equal variance. Dunnett's method for pairwise comparison was used for all post hoc sequential comparisons following one-way ANOVA; the Tukey-Kramer test was used for two-way ANOVA pairwise comparisons. Error bars in all graphs are 95% confidence intervals.

## Results

### Inoculation and infection of the brain with TMEV induces a rapid inflammatory transcriptional program

Adult female C57Bl/6 mice were intracranially inoculated with 2×10^5^ PFU of TMEV. RNA from hippocampus was prepared at 3, 6, 12, and 24 hours postinfection (hpi) and analyzed by microarray to measure changes in transcripts. Hippocampal RNA was also prepared from uninfected mice and from mice 24 hours after sham inoculation with viral growth media. Fold changes in expression for the 16900 genes present on the array were calculated relative to uninoculated/uninfected controls. A heat map of log_2_ fold change (Figure 1A) reveals that numerous genes were already upregulated at 3 hpi and the number of upregulated genes steadily grew through the first 24 hours after infection. Extraction of chemokine, cytokine, and adhesion factor genes (Figure 1B) indicates that these pathways were robustly impacted by TMEV infection. For example, the CCR2 ligand CCL2 was upregulated 8-fold by 3 hpi and continued to increase to almost 50-fold induction by 12 hpi, remaining above 30-fold at 24 hpi. The CCR5 ligand CCL5 increased steadily from 5-fold at 3 hpi to over 60-fold by 24 hpi and the CCR2 ligand CCL7 increased from 8-fold at 3 hpi to over 80-fold at 24 hpi. Likewise, the CXCR2 ligand CXCL1 was upregulated over 150-fold at 3 hpi, decreasing to 20-fold by 24 hpi, while CXCL2 was upregulated 10-fold at 3 hpi and decreased slightly to 6-fold by 24 hpi. The CX3CR1 ligand CX3CL1 was not increased at these timepoints and was, in fact, modestly downregulated by 1.4-fold at 3 hpi. In parallel with changes in chemokine expression, we observed marked upregulation of adhesion factors, with ICAM-1 upregulated 12-fold at 3 hpi, decreasing to 6-fold by 24 hpi, and VCAM-1 upregulated between 4- and 5-fold at each timepoint. Other adhesion factors involved in leukocyte trafficking were either unregulated (CD34), downregulated (Selectin-P ligand, 2-fold), or modestly upregulated (MAdCAM-1, 2-fold; GlyCAM-1, over 2-fold). Finally, a subset of immune genes were validated by RT-PCR (Figure 1C), confirming the general pattern of expression revealed by the microarray though with greater dynamic range. For example, by RT-PCR we measured a 260-fold induction of CCL2 at 3 hpi that continued to increase to over 2000-fold upregulation by 12 hpi. Overall, we conclude that inoculation of the brain with TMEV induces rapid and robust upregulation of proinflammatory chemokines, cytokines, and adhesion factors in the hippocampus that can be detected as early as 3 hours after injection.

**Figure 1.**
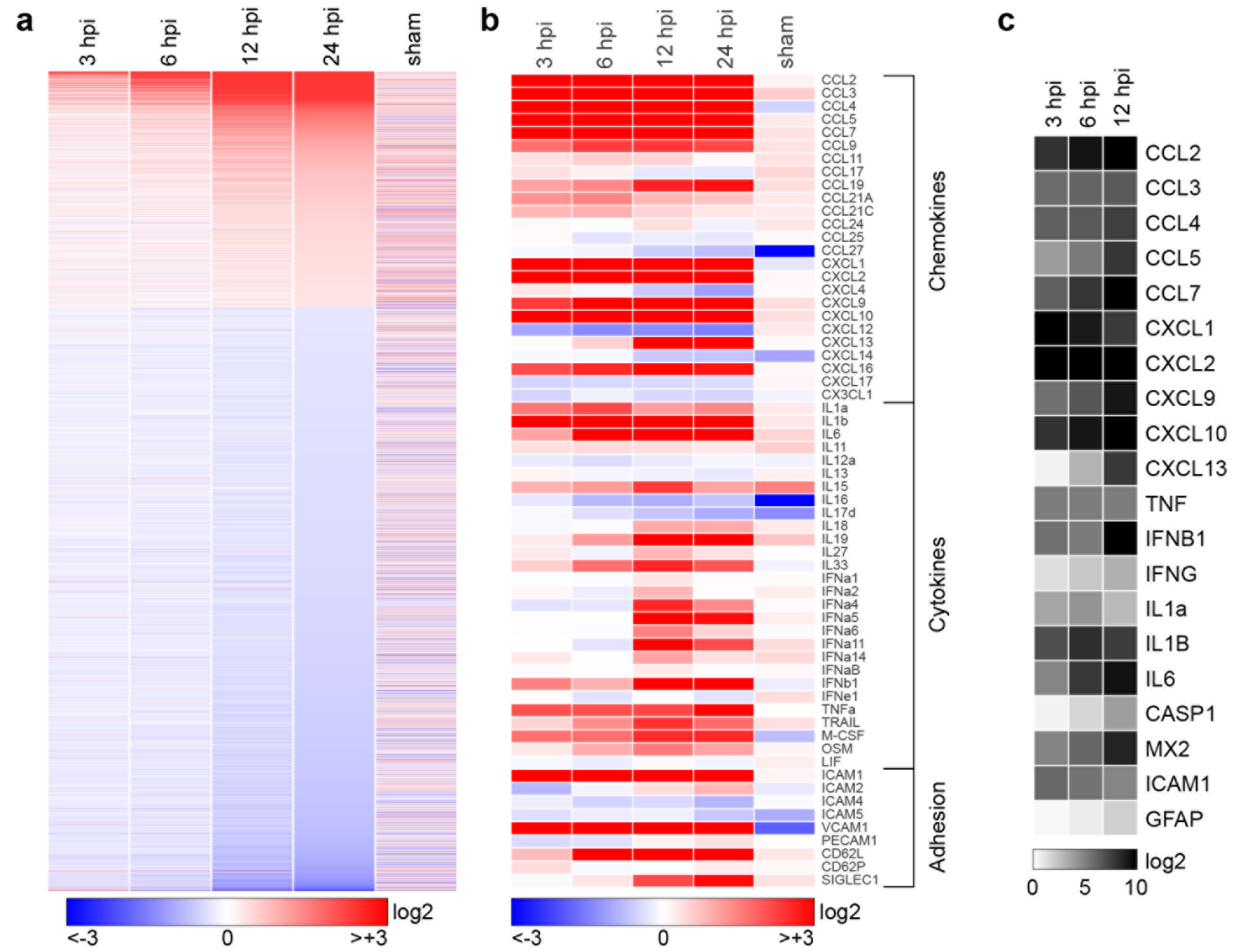
Transcriptional program initiated in the hippocampus by acute infection. Hippocampal tissue was collected for RNA purification at early timepoints following intracranial inoculation of adult female C57Bl/6 mice with 2×10^5^ PFU of the Daniel’s strain of TMEV. Transcriptional changes were analyzed using Illumina MouseWG-6 v2.0 beadchips. Fold-change relative to uninfected control for 16900 genes present on the array was calculated, converted to log_2_, and heatmapped using Gitools. Sham controls were injected intracranially with culture media used for the virus preparation and RNA was collected at 24 h postinfection (hpi). Expression data for all conditions except sham are the means from 6 individual mice analyzed in two separate experiments (3 mice per experiment). Three sham mice were analyzed in the second experiment. (A) Heatmap showing pattern of transcriptional changes at 3, 6, 12, and 24 hpi relative to uninfected controls. The discontinuity in expression coding is the result of removing from analysis all genes with changes between −1.2-fold and +1.2-fold at 24 hpi, leaving 6764 genes in the heatmap. Color-coded heatmaps range from >8-fold downregulated (log_2_<−3.0; blue) to >8-fold upregulated (log_2_>+3.0; red). (B) The same expression data remapped to highlight chemokines, cytokines, and adhesion factors. In contrast to (A), values are continuous between −8-fold and +8-fold. Note that the color range was selected to reveal weaker changes in some genes, resulting in saturation at all timepoints for highly upregulated genes such as CCL2. (C) A further subset of highly upregulated and notable genes was validated by RT-PCR and the results heatmapped on a scale from 0 to 1000-fold (log_2_=+10) to reveal large expression changes over the first 12 hours.

### TMEV infection induces rapid hippocampal production and release of the proinflammatory chemokine CCL2

Based on our previously published findings regarding hippocampal injury and inflammatory monocyte infiltration [4, 8], the known role for CCL2 in monocyte trafficking [21], and the large increase in CCL2 RNA observed as early as 3 hpi, we analyzed levels of CCL2 protein in mice acutely infected with TMEV. Serum was terminally isolated from mice inoculated with TMEV under the same conditions as above and CCL2 was measured by ELISA (Figure 2A). Circulating levels of CCL2 were detected at 300 pg/mL as early as 3 hpi, with serum levels falling by 6 and 12 hours (2A). To measure the production of this chemokine in the CNS, separate animals were perfused with PBS to remove circulating factors and whole brain was homogenized for ELISA (Figure 2B). As with the serum, CCL2 was robustly detected at 3 hpi, at levels in excess of 4000 pg per brain, and decreased sharply at 6 hpi (Figure 2B). However, in contrast to serum, the brain levels rose again at 12 hpi and reached levels comparable to the 3 hpi values by 24 hpi. Based on our previous findings regarding the locus of neuronal injury during acute TMEV infection [7, 22], we also measured CCL2 in microdissected hippocampus at the same timepoints in separate animals (Figure 2C). CCL2 levels peaked at 6 hpi and were measured at 1500 pg per mouse (hippocampi pooled for each individual animal). Notably, this amount of CCL2 represents approximately the same amount of CCL2 measured in the whole brain at this timepoint (Figure 2B), suggesting that the hippocampus is the primary site for production at 6 hpi. Finally, to separate non-infectious pathogen recognition receptor-mediated effects from chemokine induction triggered by infectious virus, especially within the context of the very rapid response following inoculation, we compared the levels of CCL2 in the whole brain at 3 hr after inoculation with UV-inactivated TMEV or live TMEV. In this experiment we measured 3010 ± 425 pg/brain with live virus, 176 ± 65 pg/brain with dead virus, and 88 ± 16 pg/brain in sham mice (live vs dead: P<0.001; live vs sham: P<0.001; dead vs sham: P=0.703; by one-way ANOVA with Dunnett's pairwise comparison). We conclude that inoculation with infectious TMEV induces the rapid production of CCL2 in the brain and particularly in the hippocampus.

**Figure 2.**
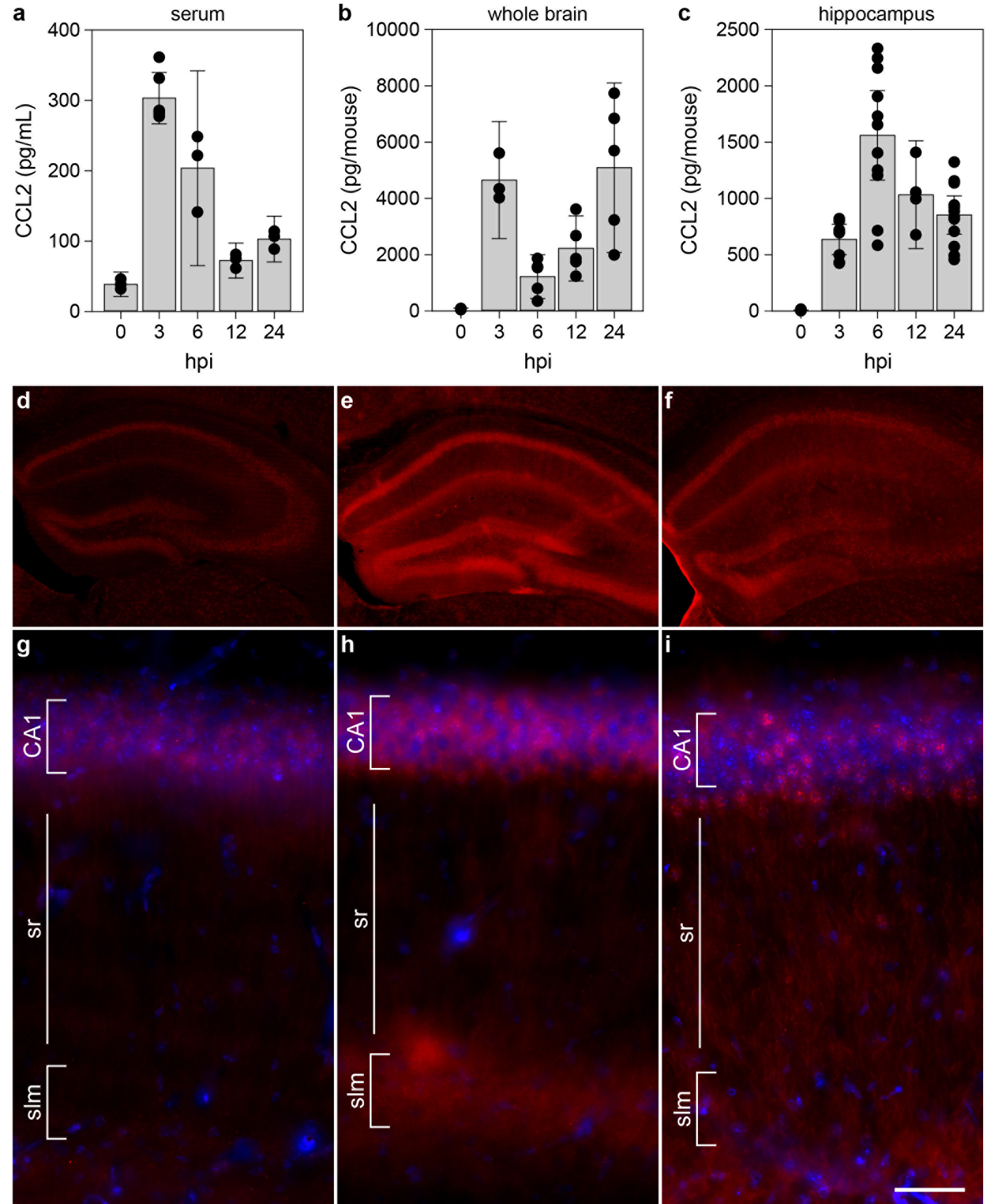
CCL2 levels in serum, whole brain, and hippocampus during acute infection. Serum (A), whole brain (B), and microdissected hippocampus (C) were collected at 0, 3, 6, 12, and 24 h following intracranial inoculation with TMEV. Absolute levels of CCL2 were measured by ELISA. Each dot represents an individual animal. The bar graph shows mean ± 95% confidence intervals from aggregate data. A minimum of 3 mice were used per tissue per timepoint and the hippocampal samples were collected in a separate experiment from the whole brain samples. All results were analyzed by one-way ANOVA. CCL2 was induced in serum, whole brain, and hippocampus (P<0.001 for each factor in each tissue compared to 0 hpi). Brain sections collected at 0 (D, G), 6 (E, H), and 24 hpi (F, I) were immunostained to reveal CCL2 expression. In the hippocampus there was weak, diffuse signal in uninfected mice (D, G) that increased markedly at 6 hpi (E, H). By 24 hpi the signal intensity had decreased and exhibited a more punctate pattern (F, I). Scale bar in I is 50 μm and refers to G-I. CA1 = Cornu Ammonis 1 formation; sr = stratum radiatum; slm = stratum lacunosum-moleculare. Immunostaining is representative of more than 3 mice per timepoint in more than 3 separate experiments.

### CCL2 is expressed in hippocampal neurons during acute infection

Given the hippocampal enrichment for CCL2 production at 6 hpi we sought to identify the relevant cellular locus (Figure 2D-I). To our surprise, immunostaining for CCL2 in uninfected mice revealed weak fluorescence associated with the pyramidal neurons of CA1, CA3, and the dentate gyrus (DG) (Figure 2D). At higher magnification this signal, not present in the secondary-only controls (not shown), exhibited a diffuse pattern associated with the cell layers (Figure 2G). At 6 hpi we consistently observed a robust increase in expression of CCL2 in pyramidal neuron cell bodies in the CA1, CA3, and DG regions (Figure 2E). In addition, there was a clear increase in immunofluorescence in the terminal networks of the apical dendrites located in the stratum lacunosum moleculare (Figure 2E, 2H). This observation is notable because we have previously reported that the hippocampal fissure, located just below the stratum lacunosum moleculare, is a primary site for infiltration of inflammatory monocytes [4]. Finally, the overall signal intensity faded by 24 hpi, though the pyramidal neuron layer-associated signal was more punctate and brighter than the earlier diffuse pattern (Figure 2F, 2I). Whether some of this signal was associated with infiltrating monocytes was not resolved in this experiment. These observations suggest that the 6 hpi peak in hippocampal CCL2 measured by ELISA (Figure 2C) was the result of production by hippocampal neurons.

### CCR2 and CCL2 are required for inflammatory monocyte infiltration during acute infection

To determine the role of the CCL2:CCR2 axis in controlling inflammatory monocyte trafficking to the brain during acute TMEV infection, we quantified leukocyte infiltration in CCR2 knockout mice at 18 hpi (Figure 3). We have previously established that our methods isolate three clearly distinguished populations that respond to TMEV infection: CD45^hi^CD11b^+^Gr1^++^1A8^−^ inflammatory monocytes, CD45^hi^CD11b^++^Gr1^+^1A8^+^ neutrophils, and CD45^mid^CD11b^mid^Gr1^−^1A8^−^ microglia [4]. In wildtype mice at 18 hpi we observed robust populations of all three cell types (Figure 3A, 3C), with inflammatory monocytes making up about 25% of the total CD45^+^ population (Figure 3E). In the absence of CCR2 expression there was marked suppression of inflammatory monocyte infiltration and a relative increase in neutrophil recruitment, with no change in microglia (Figure 3B, 3D). Indeed, inflammatory monocytes were reduced to less than 3% of total CD45^+^ cells (WT vs CCR2ko: P=0.0004 by Dunnett's method; Figure 3E). We conclude that the CCR2 receptor axis is the primary driver for inflammatory monocyte infiltration during acute TMEV infection.

**Figure 3.**
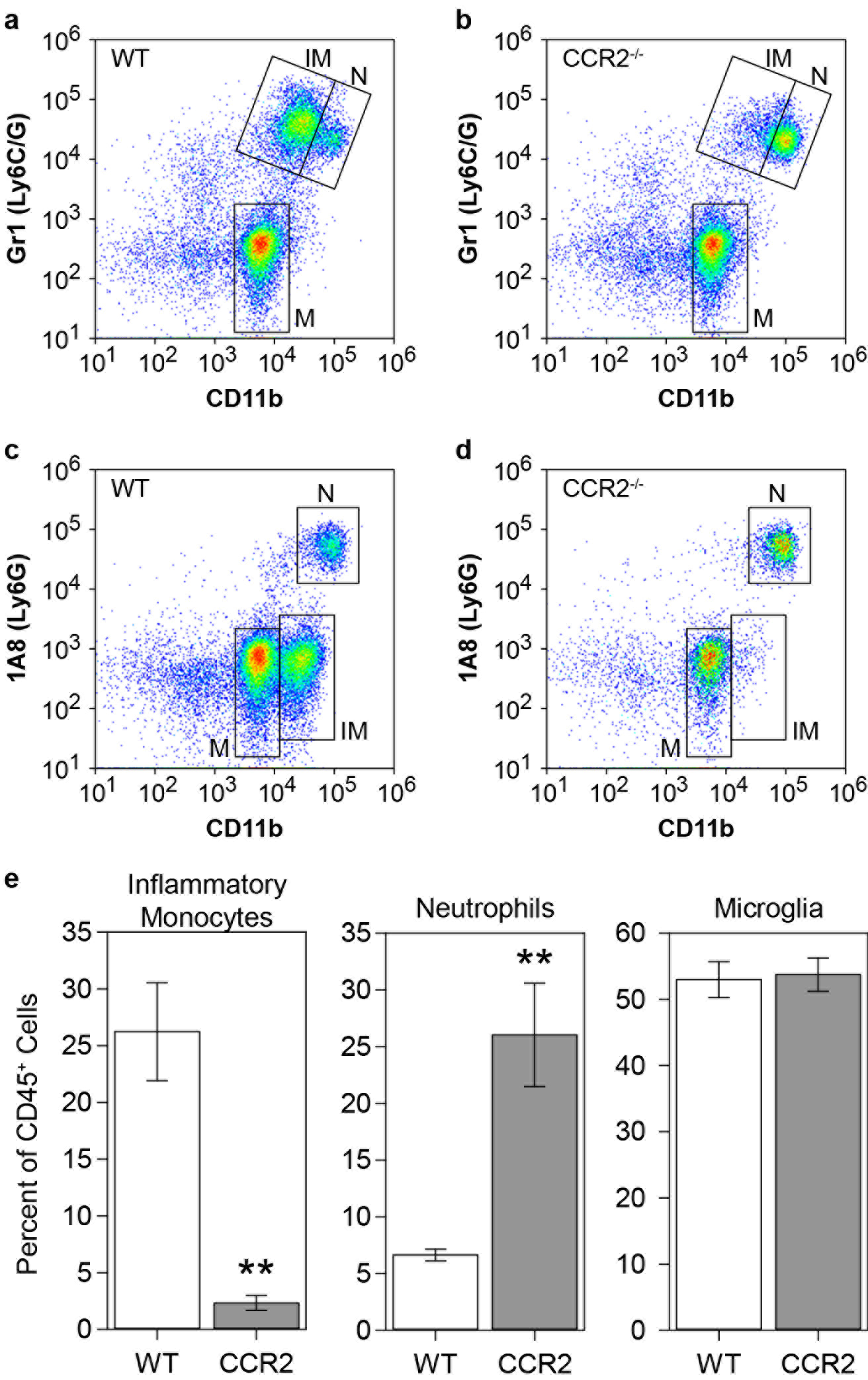
Inflammatory monocyte infiltration during acute infection requires CCR2. Brain-infiltrating leukocytes were collected from C57Bl/6 mice (WT; A, C) and CCR2^-/-^ mice (B,D) at 18 hpi. Flow plots in (A-D) show cells in a CD45^+^ parent gate. The number of inflammatory monocytes (IM; CD45^hi^CD11b^+^Gr1^++^1A8^−^), neutrophils (N;CD45^hi^CD11b^++^Gr1^+^1A8^+^), and microglia (M; CD45^mid^CD11b^mid^Gr1^−^1A8^−^) was measured by flow cytometry. The percent of inflammatory monocytes, neutrophils, and microglia in the CD45^+^ population is shown as mean ± 95%CI calculated from 9 mice per genotype in 3 separate experiments (3×3) (E). Data were analyzed by Dunnett's method. No difference in microglia was observed, but neutrophils were significantly increased in CCR2^-/-^ mice (P=0.0012 vs wt). Notably, the absence of CCR2 robustly inhibited inflammatory monocyte infiltration (P=0.0004 vs wt). **: P<0.001.

The known and putative ligands for CCR2 are CCL2, CCL7, CCL8, CCL12, CCL13, and CCL16 [23]. Of these, only CCL2 and CCL7 were expressed at detectable levels in our mice (Figure 1). To determine the relative contribution of these two chemokines to inflammatory monocyte recruitment during acute TMEV infection we used immunodepletion to neutralize each factor. LysM-GFP monocyte-neutrophil reporter mice [4] received 20 μg goat anti-CCL2 IgG, 20 μg goat anti-CCL7, or 20 μg goat IgG by intraperitoneal injection at −12, 0, and +12 hr relative to time of infection. Brain-infiltrating leukocytes were isolated at 24 hpi and analyzed by flow cytometry to count inflammatory monocytes (CD45^+^GFP^mid^) and neutrophils (CD45^+^GFP^hi^) (Figure 4). Mice treated with control IgG had approximately 5000 monocytes and 4000 neutrophils (Figure 4A, 4D) in the brain infiltrate. This was reduced to about 3500 monocytes and 3500 neutrophils in mice treated with anti-CCL7 antibody (monocytes: F=61.0887, P<0.0001 by one-way ANOVA; P<0.001 vs control IgG by Dunnett's method; neutrophils: F=54.4998, P<0.0001 by one-way ANOVA; P=0.0879 vs control IgG by Dunnett's method) (Figure 4C, 4D). More profoundly, treatment with anti-CCL2 antibody reduced the number of monocytes and the number of neutrophils to about 1500 each (monocytes: P=0.0005 vs control IgG by Dunnett's method; neutrophils: P<0.0001 vs control IgG by Dunnett's method) (Figure 4B, 4D). For monocytes, this suggests that about 70% of the infiltration in response to TMEV infection is dependent upon the CCL2:CCR2 axis and about 30% requires the CCL7:CCR2 axis.

**Figure 4.**
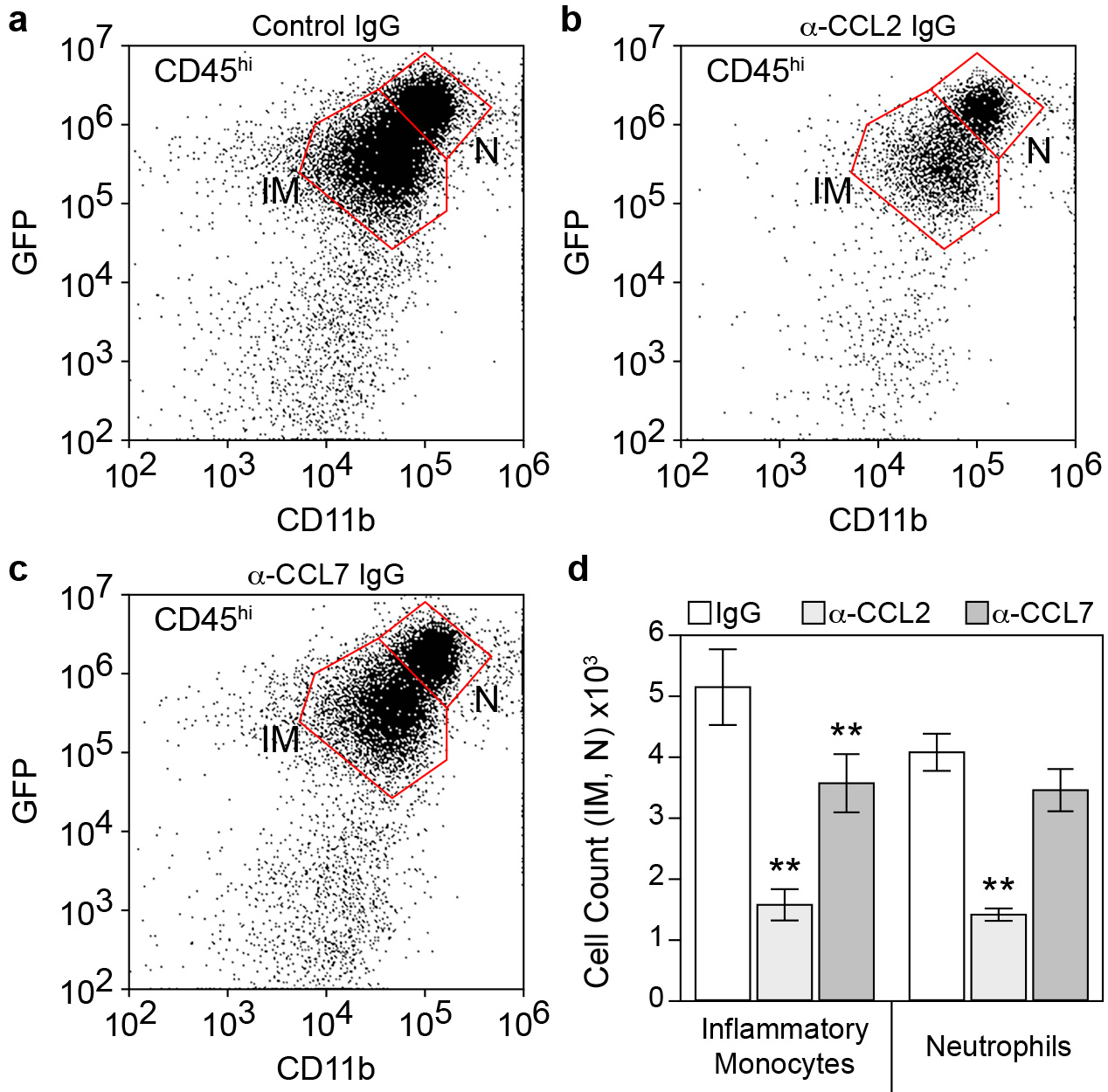
Inflammatory monocyte infiltration during acute infection is driven predominantly by CCL2. LysM:GFP mice received 20 μg goat IgG (A), 20 μg goat anti-CCL2 IgG (B), or 20 μg goat anti-CCL7 IgG (C) by intraperitoneal injection at -12, 0, and +12 hr relative to time of infection. Brain-infiltrating leukocytes were collected at 24 hpi. The flow plots in (A-C) show cells in a CD45^hi^ parent gate. The number of inflammatory monocytes (IM: CD45^hi^CD11b^+^GFP^mid^) and neutrophils (N: CD45^hi^CD11b^++^GFP^hi^) were counted; values are shown as mean ± 95% CI calculated from 6 mice per treatment condition in 2 separate experiments (3×2) (D). Data were analyzed by one-way ANOVA with Dunnett's method for pairwise comparison to control. Inflammatory monocytes were significantly reduced in both anti-chemokine groups compared to control IgG; neutrophils were significantly reduced only in the anti-CCL2 IgG group; **: P<0.001.

### Genetic deletion of CCL2 from neurons reduces serum and hippocampal CCL2 levels during acute infection

A conditional neuron-specific CCL2 knockout model was generated by crossing Ccl2-RFP^flox/flox^ mice [24] to transgenic mice expressing Cre recombinase behind the synapsin promoter (Syn-Cre) [25]. The parental Ccl2-RFP^flox/flox^ animals serve as a reporter line for CCL2 expression, and analysis of RFP fluorescence in mice at 6 hpi confirmed the CA1 neuronal expression of this chemokine (Figure 5A, 5B). Analysis of the Ccl2-RFP^flox/flox^ x Syn-Cre conditional knockouts at the same timepoint revealed that CCL2 was specifically deleted from the CA1 neurons (Figure 5C, 5D). Serum was terminally isolated from Ccl2-RFP^flox/flox^ and Ccl2-RFP^flox/flox^ x Syn-Cre mice inoculated with TMEV and CCL2 was measured by ELISA (Figure 5E). The large increase in circulating CCL2 observed at 6 hpi in Ccl2-RFP^flox/flox^ mice was not detected in the Ccl2-RFP^flox/flox^ x Syn-Cre mice (F=35.9565, P<0.0001 by two-way ANOVA; −CCL2 vs +CCL2 at 6 hpi: P<0.0001 by Tukey-Kramer pairwise analysis). However, by 24 hpi the serum levels were the same in both groups (−CCL2 vs +CCL2 at 24 hpi: P=0.9677 by Tukey-Kramer)(Figure 2E). Levels of CCL2 in the hippocampus were strongly reduced in the Ccl2-RFP^flox/flox^ x Syn-Cre mice at both 6 and 24 hpi (F=70.9728, P<0.0001 by two-way ANOVA; −CCL2 vs +CCL2 at 6 hpi: P<0.0001 by Tukey-Kramer pairwise analysis; −CCL2 vs +CCL2 at 24 hpi: P=0.0033 by Tukey-Kramer). These findings indicate that neurons are the dominant source of CCL2 in the hippocampus at 6 hpi and that neuronal CCL2 accounts for the serum levels measured at the same timepoint.

**Figure 5.**
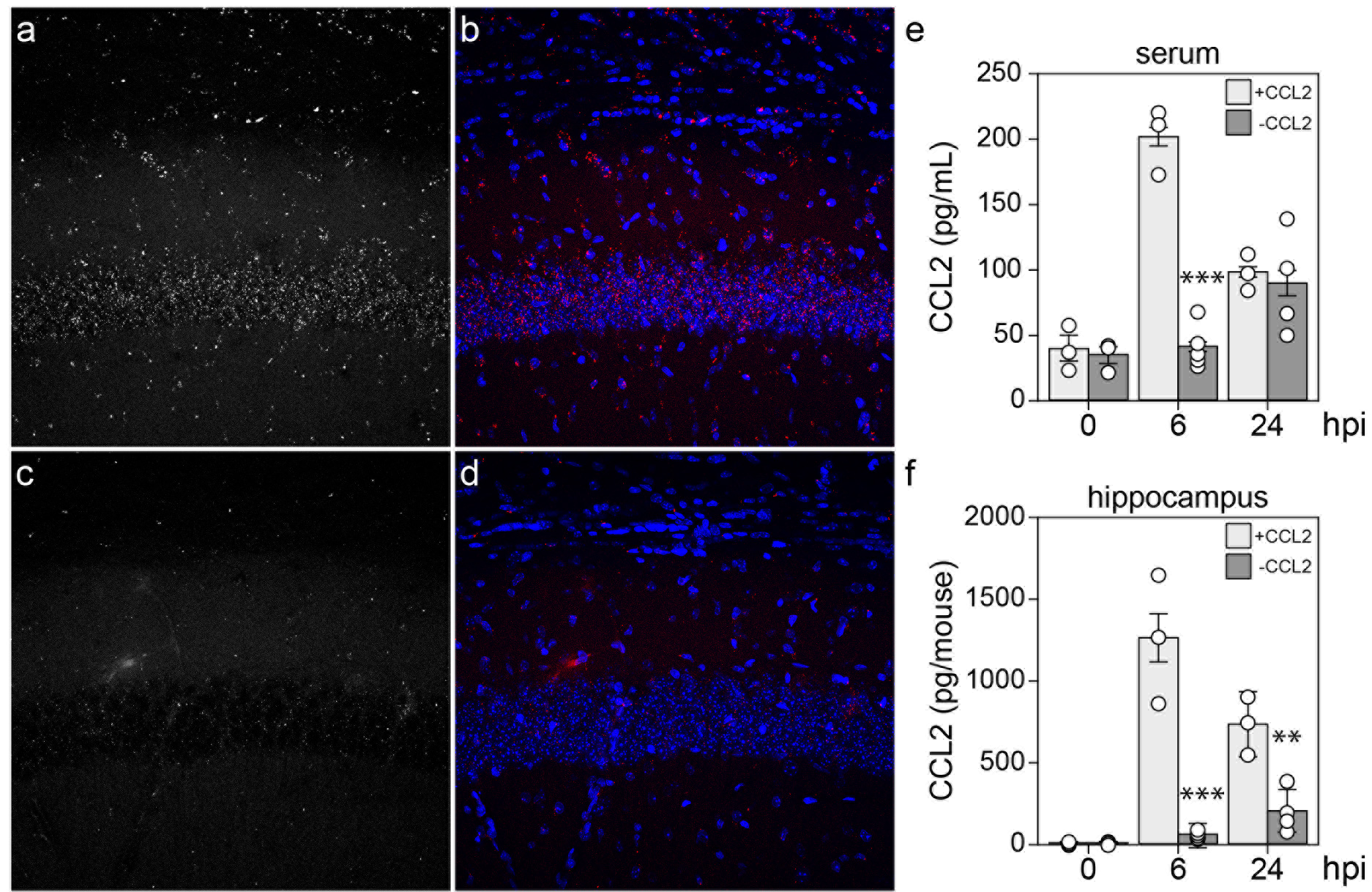
Neurons are the primary source of CCL2 at 6 hpi. Ccl2-RFP^fl/fl^ reporter mice (A, B) and Syn-Cre x Ccl2-RFP^fl/fl^ neuron-specific CCL2-deficient mice (C, D) were intracranially inoculated with TMEV and brain sections were collected at 6 hpi for analysis of RFP expression. Single-channel RFP (A, C) and two-channel RFP (red) and DAPI (blue) (B, D) microscopy revealed that the reporter signal present in neurons at 6 hpi was specifically deleted in the Syn-Cre x Ccl2-RFP^fl/fl^ mice. Serum (E) and hippocampal (F) CCL2 levels were measured by ELISA at 0, 6, and 24 hpi in the CCL2 reporter mice (+CCL2) and the neuron-specific CCL2-deficient mice (−CCL2). Each dot represents one animal; bar graphs represent mean ± 95%CI calculated from at least 3 mice per group per timepoint. Data were analyzed by two-way ANOVA with Tukey-Kramer pairwise analysis; statistical significance is only shown between genotypes at each timepoint; **: P<0.001; ***: P<0.0001.

### Genetic deletion of CCL2 from neurons reduces inflammatory monocyte infiltration during acute infection

The profound reduction in hippocampal CCL2 at 6 hpi in the neuron-specific CCL2-deficient mice suggested that monocyte infiltration would also be impaired in these animals. Brain-infiltrating leukocytes were collected from wildtype B6 mice (Figure 6A), CCL2-RFP^fl/fl^ reporter mice (+CCL2; Figure 6B), and CCL2-RFP^fl/fl^ x Syn-Cre mice (−CCL2; Figure 6C) at 18 hpi and analyzed by flow cytometry. Gates were established as described above on a CD45^hi^ parent gate. The total number of CD45^hi^ cells isolated from B6 mice and CCL2-RFP^fl/fl^ reporters was not different (F=44.8489, P<0.0001 by one-way ANOVA; B6 vs +CCL2: P=1.000 by Dunnett's method) but the number of these cells in the neuron-specific CCL2-deficient mice was significantly reduced (−CCL2 vs B6: P=0.0001; −CCL2 vs +CCL2: P=0.0004) (Figure 6D). The trend toward more CD45^hi^ cells in the CCL2 reporter mice suggested by Figure 6D was exacerbated in the neutrophil counts (Figure 6E), though the difference was not significant (F=87.2340, P<0.0001 by one-way ANOVA; B6 vs +CCL2: P=0.1908 by Dunnett's method). In contrast, the number of neutrophils infiltrating the brain was reduced in the neuron-specific CCL2-deficient mice (−CCL2 vs B6: P=0.0003; −CCL2 vs +CCL2: P<0.0001) (Figure 6E). Finally, the number of inflammatory monocytes in the brain at 18 hpi did not differ between B6 and reporter mice (F=56.8592, P<0.0001 by one-way ANOVA; B6 vs +CCL2: P=1.000 by Dunnett's method) but was significantly reduced in the neuron-specific CCL2-deficient mice (−CCL2 vs B6: P<0.0001; −CCL2 vs +CCL2: P=0.0024) (Figure 6F). These observations support the conclusion that the limited CCL2 production and release measured in neuron-specific CCL2-deficient mice at 6 hpi results in a strong reduction in inflammatory monocyte infiltration into the brain at 18 hpi.

**Figure 6.**
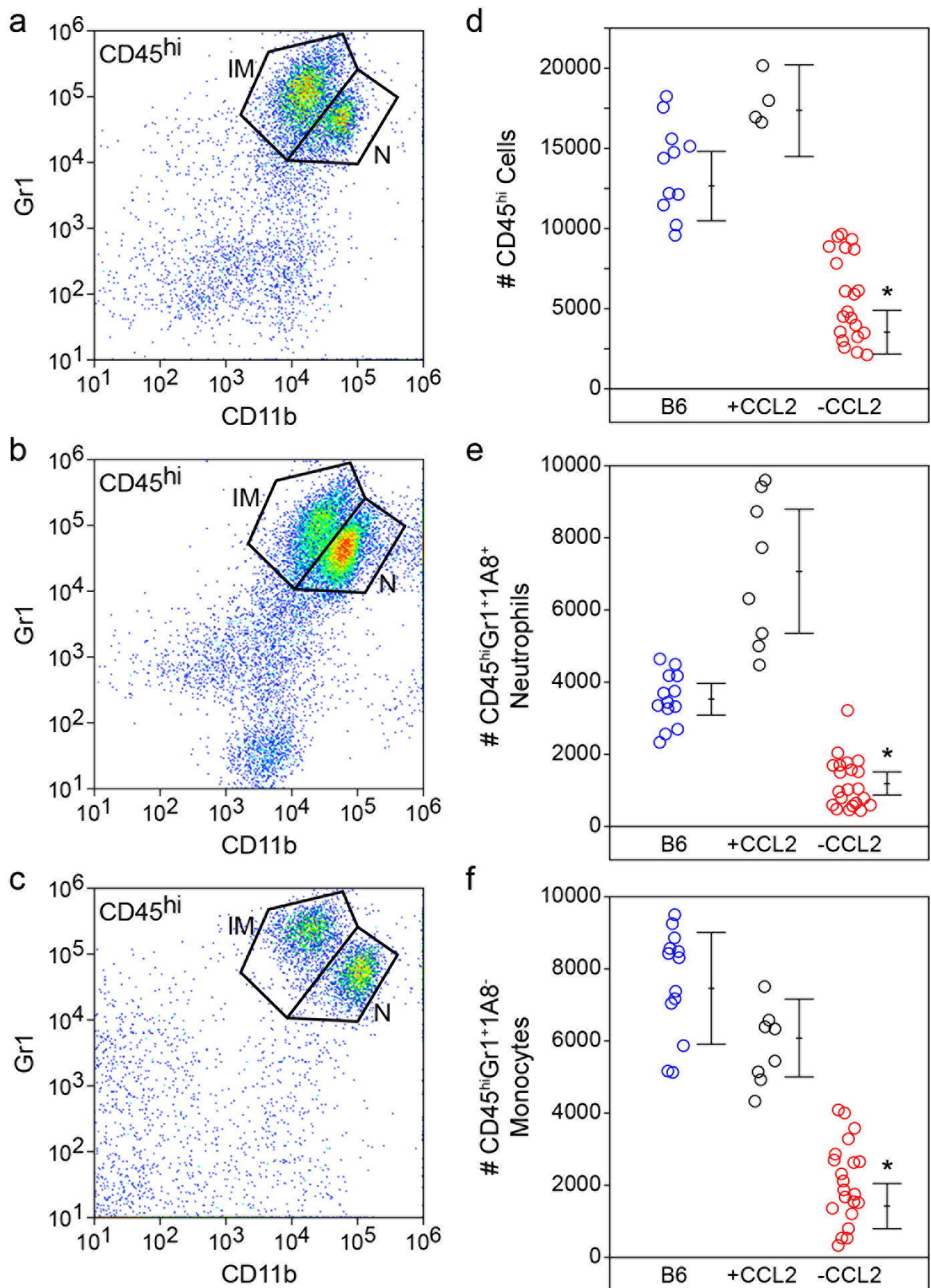
Loss of acute neuronal CCL2 production results in reduced inflammatory monocyte infiltration into the brain. Brain-infiltrating leukocytes were analyzed by flow cytometry in wildtype B6 mice (A), Ccl2-RFP^fl/fl^ reporter mice (^+^CCL2; B), and Syn-Cre x Ccl2-RFP^fl/fl^ neuron-specific CCL2-deficient mice (−CCL2; C) at 18 hpi. The flow plots in (A-C) show Gr1- and CD11b-labeled cells in a CD45^hi^ parent gate. The number of CD45^hi^ cells (D), neutrophils (E), and inflammatory monocytes (F) were counted in the three groups (blue circles = B6; black circles = +CCL2; red circles = −CCL2). Each dot represents one animal; the line graph represents mean ± 95%CI calculated from at least two separate experiments. All cell types were significantly reduced in the neuron-specific CCL2-deficient mice but not in the parent reporter line. Data were analyzed by one-way ANOVA with Dunnett's pairwise comparison; *: P<0.0001.

## Discussion

Identification of immune cells and effector molecules that contribute to brain injury is critical to the development of neuroprotective and neuroreparative strategies [7]. We and others have used the Theiler's murine encephalomyelitis virus model to investigate immune-mediated mechanisms of brain injury within the context of acute CNS viral infection [3, 6]. Despite differences in nomenclature, host genetics, and details of the viral strain and inoculum, a general consensus has emerged that monocytic cells infiltrate the brain within hours of infection and create an inflammatory environment that kills neurons and alters neural circuitry. Understanding the mechanisms that drive monocyte infiltration may therefore reveal novel strategies to protect the brain.

Not surprisingly, intracranial inoculation with TMEV induced a robust inflammatory program in the hippocampus, involving upregulation of dozens of chemokine, cytokine, and adhesion factor genes (Figure 1). Somewhat surprising, however, was the speed with which this induction occurred. We reproducibly detected large changes in transcription by 3 hpi, and in some experiments measured variable, but sizable, induction of chemokines and cytokines by 1 hpi (not shown). Moreover, these factors were detected at the protein level over the same acute period, with CCL2 measurable in the serum and brain as early as 1 hpi (not shown). Not only are these responses fast with regard to biosynthesis, but it is unlikely that a significant number of cells have even been infected in the brain by 3 hpi. Mathematical modeling constrains the infection rate to only 0.01 to 0.1 cells per minute [26], suggesting that fewer than 20 cells would be productively infected by 3 hpi. Even if this rate is off by a factor of 100, fewer than 2000 cells are infected by 3 hpi. Obviously, some cells can respond to virions and virus constituents via mechanisms that do not require active infection, such as pathogen (or pattern) recognition receptors [27], but our findings indicate that inoculation with UV-inactivated virus did not elicit CCL2 production in the brain at 3 hpi. This finding also rules out induction via non-specific trauma-induced effects associated with intracranial inoculation. Moreover, even assuming a direct effect of each active virion on target cells, we only inoculated the animals with 200,000 plaque forming units and these were not introduced directly into the hippocampus [20]. Yet, at 6 hpi the hippocampus produced essentially all of the CCL2 measured in the brain and presumably contributed the majority of serum CCL2. This issue is further confounded by the pattern of CCL2 expression revealed by immunostaining, which shows that effectively every neuron in dorsal hippocampal CA1, CA3, and dentate gyrus expressed this factor at 6 hpi (Figure 2). As we have previously reported, even by 3 dpi only a fraction of CA1 neurons are directly infected with TMEV and DG neurons are never positive for virus by immunostaining [22]. These observations and discrepancies suggest that a currently unidentified amplification event occurs almost immediately after inoculation with live virus that results in widespread, albeit tissue-specific, upregulation of CCL2 production. Furthermore, this induction occurs almost exclusively in neurons, as synapsin-promoter driven deletion of CCL2 results in nearly complete suppression of both hippocampal and serum CCL2 at 6 hpi.

Despite the gap in our knowledge about the pathway between virus inoculation and CCL2 induction, our data clearly support a model in which neurons are the primary source of this chemokine during the most acute phase of the host response. Notably, deletion of CCL2 from neurons recapitulates the effect of systemic deletion in mice treated with anti-CCL2 immunoglobulin (Figure 6F compared to Figure 4D) - the inflammatory monocyte infiltrate is reduced by about 70% in both conditions at 18-24 hpi. Presumably, the remaining infiltrate in both experiments is CCL7-dependent, although the cellular source of this other CCR2 ligand is currently unknown. These findings suggest an intriguing maladaptive response: neurons respond to brain inoculation with TMEV by producing copious amounts of a chemokine that serves to recruit the inflammatory monocytes into the brain that ultimately kill those same neurons [4]. While the current study does not address the impact of a curtailed monocyte response on viral clearance or eventual lymphocyte recruitment, anecdotal evidence indicates that CCR2^-/-^ mice do not succumb to lethal infection and, indeed, do not show any apparent adverse effects at later timepoints (out to several months). Studies assessing the impact of CCR2 deletion on hippocampal neuropathology are ongoing.

Maintaining the focus on the most acute response to TMEV infection rather than downstream sequelae, several notable findings arise from the current study. First, genetic deletion of CCR2 in all cells almost entirely abrogates inflammatory monocyte infiltration at 18 hpi but actually increases neutrophil infiltration at this timepoint. This suggests that in the context of life-long absence of CCL2:CCR2 signaling neutrophil responses are not compromised by the absence of an inflammatory monocyte response. Second, acute systemic immunodepletion of CCL2 reduces inflammatory monocyte infiltration by 70% but also reduces neutrophil infiltration by more than half. This suggests that acute inhibition of the CCL2:CCR2 axis either impacts neutrophil recruitment directly (for example, brain infiltrating neutrophils express CCR2 and respond to CCL2) or via an indirect effect on monocyte-to-neutrophil communication (for example, infiltrating monocytes release neutrophilic chemokines) [28]. Third, systemic immunodepletion of CCL7 blocks about 30% of monocyte infiltration but has no impact on neutrophils. This indicates that if brain infiltrating neutrophils use a CCR2-dependent trafficking pathway it is not responsive to CCL7 or that the CCL7-dependent brain-infiltrating monocytes are not responsible for creating the pro-neutrophil environment. While speculative, this may suggest that there are at least two functional subtypes of CCR2^+^ brain-infiltrating monocytes that can be distinguished by CCL2 vs CCL7 responsiveness. Fourth, the CCL2-RFP^fl/fl^ reporter line exhibits a trend toward increased neutrophil infiltration as compared to B6 mice. While not statistically significant due to the spread in values, this increase was visually evident in most of the mice, suggesting caution in studies analyzing leukocytic responses in these animals. Fifth, the robust inhibition of monocyte infiltration in the neuron-specific CCL2-deficient mice argues explicitly that this chemokine drives trafficking to the brain while side-stepping all of the issues regarding monocytopenia that arise in CCL2^-/-^ or CCR2^-/-^ mice. This is in contrast to the reported role of CCR2 in West Nile virus encephalitis [14] but is consistent with the brain trafficking role described by Graham and colleagues in a Semliki Forest virus model [29]. Sixth, manipulation of either CCL2 or CCR2 drove the inflammatory monocyte response in the same direction - down. This is in contrast to observations in a Japanese encephalitis model, in which CCR2^-/-^ mice had reduced monocyte infiltration while CCL2^-/-^ mice had a paradoxical increase in this population in the brain [16]. Of note, however, our manipulation of CCL2 was via acute immunodepletion, not life-long genetic deletion.

The production of CCL2 by neurons during acute encephalitis may play a role in more than just leukocyte recruitment. For example, hippocampal neurons express CCR2 [21] and CCL2 directly enhances both NMDA receptor- and AMPA receptor-mediated excitatory postsynaptic potentials [30]. This suggests that the earliest stages of hippocampal circuit dysregulation associated with acute TMEV infection may be triggered locally by neuronal CCL2 acting in a autocrine fashion [31, 32]. In this context, it is notable that CCL2 immunoreactivity is strongly upregulated in CA1 neuron apical dendrite tufts located in the stratum lacunosum moleculare at 6 hr after infection (Figure 2). This layer is an important site of integration between the entorhinal cortex and CA1 pyramidal neurons (the perforant pathway) and is of fundamental importance to spatial and episodic memory formation [33]. It is also the site of GABAergic interneurons that are strongly activated by both entorhinal cortex and CA3-derived Schaffer collateral inputs [34]. These interneurons exhibit both AMPA receptor and NMDA receptor excitatory postsynaptic potentials [35] and are maximally activated at the theta oscillation peak and at the gamma oscillation trough [36]. These cells appear to mediate a robust feedforward inhibition that controls the size and timing of excitatory inputs onto CA1 neurons [37] and synchronizes network firing to theta frequency [34, 36]. Given the important role for hippocampal theta oscillations in learning and memory [38], it is likely that acute production of CCL2 in the stratum lacunosum moleculare would disrupt cognitive performance. It is also notable that neuronal CCL2 production has been observed following kainic acid- [39] and pilocarpine-mediated seizure induction [40] in rodents and both CCL2 and CCR2 are increased in tissue resected from humans with intractable epilepsy [41, 42]. Inhibition of either CCL2 production or CCR2 signaling suppressed seizures in a mouse model of systemic inflammation and mesial temporal lobe epilepsy [43]. Indeed, Caleo and colleagues have recently suggested that CCL2 "may act as a master regulator of inflammatory processes in the epileptic brain by both directly promoting hyperexcitability and regulating the activity of downstream inflammatory effectors" [43]. Our findings echo this concept and add a third role in which hyperacute neuronal CCL2 production mediates CCR2-dependent inflammatory monocyte trafficking into the brain, resulting in the creation of an environment that further disrupts neural function and induces pyramidal neuron death [4, 7].

## Conclusion

Overall, we conclude that infection of the brain with TMEV rapidly induces CCL2 expression in neurons and this cellular source is central to driving CCR2-dependent infiltration of inflammatory monocytes into the brain during the most acute stage of encephalitis. While the relevance of our current findings to the neuropathological and electrophysiological sequelae of TMEV encephalitis remains to be determined, our observations lend further support to the therapeutic relevance of targeting the CCL2:CCR2 axis to confer neuroprotection. Our findings also highlight a unique role for neuronal production of chemokines in the initiation of leukocytic infiltration into the infected central nervous system.

## Abbreviations

AMPA: α-amino-3-hydroxy-5-methyl-4-isoxazolepropionic acid
ANOVA: analysis of variance
BILs: brain-infiltrating leukocytes
BSA: bovine serum albumin
hpi: hours postinfection
CCL: CC motif chemokine ligand
CCR: C-C motif chemokine receptor
CNS: central nervous system
CXCL: C-X-C motif chemokine ligand
CXCR: C-X-C motif chemokine receptor
CX3CL: C-X3-C motif chemokine ligand
CX3CR: C-X3-C motif chemokine receptor
DAPI: 4′,6-diamidino-2- phenylindole
DMEM: Dulbecco’s modified Eagle's medium
ELISA: enzyme-linked immunosorbent assay
FBS: fetal bovine serum
GABA: gamma-aminobutyric acid
GFP: green fluorescent protein
GlyCAM: glycosylation-dependent cell adhesion molecule
ICAM: intercellular adhesion molecule
KO: knockout
MAdCAM: mucosal vascular addressin cell adhesion molecule
NMDA: N-methyl-D-aspartate
PFA: paraformaldehyde
PFU: plaque-forming units
PBS: phosphate-buffered saline
RFP: red fluorescent protein
RT-PCR: reverse transcription polymerase chain reaction
TMEV: Theiler's murine encephalomyelitis virus
UVc: ultraviolet C
VCAM: vascular cell adhesion molecule
WT: wildtype

## Acknowledgements

We thank the Mayo Medical Facility Gene Expression Core for assistance with microarray analysis. We thank Dr. Ben Clarkson, Renee Johnson, and Misha Patel for their expert technical assistance. We thank Dr. Shai Kunda, Erin Triplet, and Ethan Grund for their input on the manuscript.

## Funding

This work was supported by funding from the NINDS/NIH to CLH (NS064571) and by postdoctoral funding from the Mayo Clinic Center for Multiple Sclerosis and Autoimmune Neurology to KM. ENG was supported by the Mayo Clinic Regenerative Sciences PhD training program.

## Authors’ contributions

CLH conceived the experiments, established the experimental design, analyzed all of the data, wrote the manuscript, and prepared the final figures. RLC, ENG, and KM performed experiments, analyzed the data, and contributed to the figures; RLC and ENG provided text and edited the manuscript. All authors read and approved the final manuscript.

## Ethics approval

All animal experiments were performed according to the National Institutes of Health guidelines and were approved by the Mayo Clinic Institutional Animal Care and Use Committee (Animal Welfare Assurance number A3291-01).

## Competing interests

The authors declare no competing interests.

